# Enhanced anticipatory biasing of visuospatial attention in deaf native-signing adults indexed by alpha-band (8-14 Hz) oscillatory neural activity

**DOI:** 10.1101/2022.01.25.477746

**Authors:** Ian A. DeAndrea-Lazarus, Edward G. Freedman, Jiayi Xu, Kevin D. Prinsloo, Maeve M. Sargeant, John J. Foxe

**Author notes:** Correspondence: John J. Foxe, Ph.D., Department of Neuroscience, University of Rochester Medical Center, 601 Elmwood Avenue - Box 603, Rochester, New York 14642.

## Abstract

Deaf people show increased visuospatial attention abilities, especially towards peripheral inputs, but the neural mechanisms of these heightened abilities are not yet understood. In hearing individuals, topographically-specific alpha-band oscillatory activity (8-14 Hz) over parieto-occipital regions has been associated with active suppression of irrelevant locations. Here, we asked whether increases in this spatially-specific anticipatory oscillatory mechanism might underpin enhanced visuospatial attention abilities in deaf individuals, on the premise that deaf people might be more adept at transiently engaging and disengaging attentional processes involved in processing peripheral inputs. An alternative hypothesis was that deaf individuals might not produce lateralized alpha-band activity, because of the need to continuously monitor the periphery due to the absence of an auxiliary auditory spatial alerting system. High-density electroencephalography was recorded from 20 deaf native signers and 20 hearing non-signers performing a cued covert visuospatial attention task. Deaf participants responded significantly more rapidly and accurately and showed highly typical alpha-band lateralization during the cue-target interval of the task. Topographic analysis showed a greater extent of alpha-band anticipatory activity over right parietal scalp, suggesting sequestration of extra-visual attentional circuits (i.e., unused auditory regions), and *post-hoc* analysis pointed to substantially earlier onset of this activity during the cue-target interval. The presence of cue-evoked anticipatory alpha lateralization in deaf participants suggests that they are rapidly engaging and disengaging attentional processes involved in orienting attention to the periphery. The earlier and more extensive engagement of these anticipatory oscillatory processes may contribute to the improved visuospatial performance observed in these individuals.

**Significance Statement:** Prior to this study, it was not known whether deaf people demonstrate lateralization of alpha-band oscillatory electroencephalographic (EEG) activity over the posterior region of the brain, which plays a role in the suppression of uncued regions of space during cued visuospatial attention tasks. We found that this lateralized pattern was observable in deaf participants and was not significantly different from that seen in hearing participants, except that alpha activity onsets earlier in deaf participants. However, when cue directions were collapsed, the scalp topographies of deaf participants showed a greater distribution of alpha activity, suggesting that they recruited a brain region typically reserved for audiospatial attentional control during the visuospatial attention task. Additionally, deaf participants responded significantly more quickly and accurately compared to hearing participants, demonstrating increased visuospatial attention abilities.

## Introduction

Given limited access to sound, deaf and hard-of-hearing people are more reliant on vision for environmental cues than their hearing counterparts, who also have the benefit of their auxiliary auditory spatial alerting system. As a result, deaf people may develop enhanced compensatory visuospatial attentional skills, especially for inputs to the periphery. Consistent with this notion, deaf individuals are faster at detecting and discriminating peripheral stimuli without sacrificing accuracy (Parasnis and Samar, 1985; Neville and Lawson, 1987a; Loke and Song, 1991; Reynolds, 1993; Colmenero et al., 2004; Chen et al., 2006; Nava et al., 2008; Dye et al., 2009; Bottari et al., 2010, 2011), findings that are in line with the sensory compensation hypothesis, which posits that “*a deficit in one sensory system would make other modalities more sensitive, vicariously compensating for the loss of one sensory channel*” (Pavani and Bottari, 2012). One possible explanation for increased sensitivity to peripheral distractors (Bosworth and Dobkins, 2002; Proksch and Bavelier, 2002; Rothpletz et al., 2003) is that deaf individuals may constitutively sustain heightened spatial attention in readiness for potential peripheral inputs as an early warning system, a potentially costly neural strategy (Laughlin et al., 1998). Alternately, increased peripheral processing might be driven by increased facility in the transient deployment of voluntary (endogenous) spatial attentional processes, allowing for rapid online engagement and disengagement from to-be-attended locations (Posner et al., 1980; Parasnis and Samar, 1985; Bosworth and Dobkins, 2002; Lalor et al., 2007).

A common approach to assessing the ability to engage and disengage peripheral attentional processes is to employ a cued covert visuospatial attention task (Posner, 1980). In response to a directional cue, attentional resources are voluntarily deployed to the cued location where stimuli are expected to occur without explicitly moving the direction of one’s gaze to that location (Posner, 1980; Hillyard and Anllo-Vento, 1998; Simpson et al., 2006; Kelly et al., 2008). Neurophysiological recordings allow for the evaluation of attentional processing during the cue-target interval and provide insights into the transient engagement of attentional biasing processes that are deployed in anticipation of upcoming targets (and distractors) (Foxe et al., 2005; Dale et al., 2008; Van Diepen et al., 2019).

Spatially-specific power increases (and decreases) in the alpha-oscillatory band (∼8-14 Hz) have been repeatedly shown to be a robust index of the anticipatory deployment of visuospatial attention during the cue-target interval of a visuospatial cueing task (Worden et al., 2000; Yamagishi et al., 2003; Kelly et al., 2005, 2006, 2009; Sauseng et al., 2005; Rihs et al., 2007, 2009). Scalp topographies are consistent with receptive fields matching the location of impending distractors, lending support to the alpha suppression hypothesis, which posits that these increases in posterior alpha-band activity reflect attentional suppression mechanisms (Foxe et al., 1998; Kelly et al., 2006; Thut et al., 2006; Klimesch et al., 2007; Jensen and Mazaheri, 2010; Foxe and Snyder, 2011; Belyusar et al., 2013; Gray et al., 2015; Foster and Awh, 2019; Sokoliuk et al., 2019; Van Diepen et al., 2019; Wöstmann et al., 2019; Hutchinson et al., 2020; Wilson and Foxe, 2020). Based on previous findings, we along with others have suggested that the spatially specialized dorsal visual stream in the posterior parietal cortex exerts control over the gateways to the ventral visual stream, which is specialized for processing featural information (Mishkin et al., 1983; Dockree et al., 2007; Foxe and Snyder, 2011; Capilla et al., 2014; Zumer et al., 2014; Zhigalov and Jensen, 2020). Thus, alpha activity in the posterior parietal cortex (PPC) may represent a gating mechanism in which an increase in alpha activity represents a selective inhibition of featural processing of irrelevant stimuli.

There is evidence of compensatory plastic changes in deaf native signers as a result of limited auditory access, especially in brain regions associated with visuospatial processing (Neville et al., 1983; Neville and Lawson, 1987a; Loke and Song, 1991; Bavelier et al., 2001; Pavani and Bottari, 2012; Dye and Bavelier, 2013). A deaf native signer is defined as a prelingually deaf person, born to deaf parents, who has acquired sign language as their first language during the typical timeframe for language development (Dye and Bavelier, 2013). Of note, there is greater recruitment of the posterior parietal cortex (PPC) and enhanced functional connectivity between the PPC and earlier visual areas in deaf signers compared to hearing participants as well as hearing native signers when processing peripheral stimuli (Bavelier et al., 2000, 2001; Scott et al., 2014). Here we asked whether potential heavier reliance on peripheral vision by deaf individuals would lead to enhanced visuospatial attention abilities, reflected by increases in spatially specific anticipatory alpha-band oscillatory processes. In this case, we theorized that deaf individuals might be more adept at transiently engaging and disengaging processes involved in the processing of potential inputs of interest or distractors outside the focus of attention. An alternative hypothesis, however, is that deaf individuals might not engage this selective suppression mechanism as strongly as hearing individuals due to a need to maintain consistent visual monitoring for potentially relevant (or threatening) peripheral inputs as a result of not possessing an auxiliary auditory spatial alerting system. This alternate hypothesis derives some support from experimental evidence that deaf participants are more easily distracted by irrelevant peripheral stimuli (Proksch and Bavelier, 2002; Sladen et al., 2005; Dye et al., 2007).

A second major focus of this study pertains to potential sensory-substitution processes in the overall control of attention in deaf individuals. Earlier work by our group has demonstrated that there is a supramodal spatial attention system as well as sensory-specific control systems in the right parietal lobe of hearing individuals (Banerjee et al., 2011). Hearing participants engaged in separate visuospatial and audiospatial cueing tasks initially displayed a highly focused right parieto-occipital alpha distribution shared by both modalities and then displayed distinct auditory and visual alpha foci over the parietal scalp in the late anticipatory period independent of cue direction. This differential distribution for each modality indicates that there are sensory-specific control systems within the right parietal attention control system (see also, Krumbholz et al., 2009). In the present study, we were interested in understanding the implications for right parietal control of spatial attention if one of the sensory systems represented in this system is unused. We hypothesize that the topography of the right parietal alpha-band control process would have greater extent in deaf participants relative to hearing participants, reflecting potential sequestration of auditory spatial control circuitry by the visuospatial attentional system in deaf individuals.

## Materials and Methods

### Participants

Twenty-one deaf adults (14 female), aged 28.0 ± 5.3 years, and twenty hearing adults (12 female), aged 22.1 ± 3.6, were enrolled. All participants reported no history of neurological or neurodevelopmental disorders, did not have a history of head injury, were not dependent on drugs or alcohol, were not under the influence of drugs or alcohol at the time of the study, and had normal or corrected-to-normal vision. All deaf participants were native signers – they were born deaf to deaf parents and acquired American Sign Language (ASL) as their first language. All deaf participants reported bilateral hearing loss and the average hearing loss in the better ear was 91 ± 12 decibels. All hearing participants reported no previous knowledge of ASL. All participants provided written informed consent, and the Research Subjects Review Board of the University of Rochester reviewed and approved all experimental procedures. This study conforms to the principles outlined in the Declaration of Helsinki. Participants received a modest fee of $15 per hour for their participation.

### Stimuli and task

Participants were seated in an electromagnetically shielded and acoustically dampened booth and rested their chin comfortably on an adjustable chinrest 0.8 m from a curved computer monitor (Acer Predator Z35). Visual stimuli were presented on a black background using Presentation (Neurobehavioral Systems, Albany, CA). The cued visuospatial attention task used in this study was adapted from Vollebregt et al. (2015) and is shown in Figure 1. Participants were shown an instructional video in either spoken English or American Sign Language before starting the task. At the beginning of each trial, participants were shown a static image of a cartoon fish with edges subtending 1.16 degrees from the midline (2.32 degrees from edge to edge), looking back at the participant from the center of the screen. There were also two static images of the profile of a shark positioned on each side of the central fish on the horizontal plane such that it appears that the two sharks are looking directly at the fish. The visual angle for the innermost edge of the two sharks was 14.5 degrees from the center of the screen. The size of the shark was 7 degrees on the horizontal plane, and the most informative part of the shark was the inner half where the mouth was located (14.5 to 18 degrees). Participants were instructed to fixate on the central fish for the duration of the task. An EyeLink 1000 infrared eye tracker (SR Research) was used to ensure fixation at all times. Nine-point calibration of the eye tracker was performed at the beginning of the experiment and after the mid-task break. After 500 ms in neutral gaze position (eyes open and looking straight ahead), the central fish would briefly shift its gaze left or right towards either shark for 100 ms (the spatial cue). After a 1000 ms interval, one of the sharks (the target) would open its mouth wider for 100 ms. The participant then had up to 1400 ms to press either the left or right response button on a standard keyboard with their right index finger or right middle finger, respectively. A cue validity of 50% was used, and targets had an equiprobable chance of appearing in either hemifield. Participants were instructed to press the button corresponding to the direction of the gaze cue only. That is, the task was to deploy covert spatial attention to the side indicated by the cue and only respond to targets appearing there. Those targets appearing in the uncued hemifield were to be ignored. The need to withhold response for these invalid cues ensured that the cue remained informative even though it was only valid 50% of the time. After a response was made or if time expired, the participant was shown a feedback image centrally. This image was either a positive (fish flapping its fins) or a negative (fish skeleton) feedback token. The task consisted of 368 trials evenly divided over two blocks with five brief breaks within each block, during which a short clip (7 ± 2 seconds) giving positive reinforcement was shown (i.e., praise for a job well done). If participants’ gaze deviated from central fixation or if there was excessive blinking, the trial was aborted, and a 7-second video clip was shown reminding the participant to fixate on the central cue. Aborted trials were added back into the trial block to ensure equal numbers of trials. One deaf participant did not understand the task and their data were contaminated with button presses during the cue-target interval, so their data was not included in the analyses.

**Figure 1.**
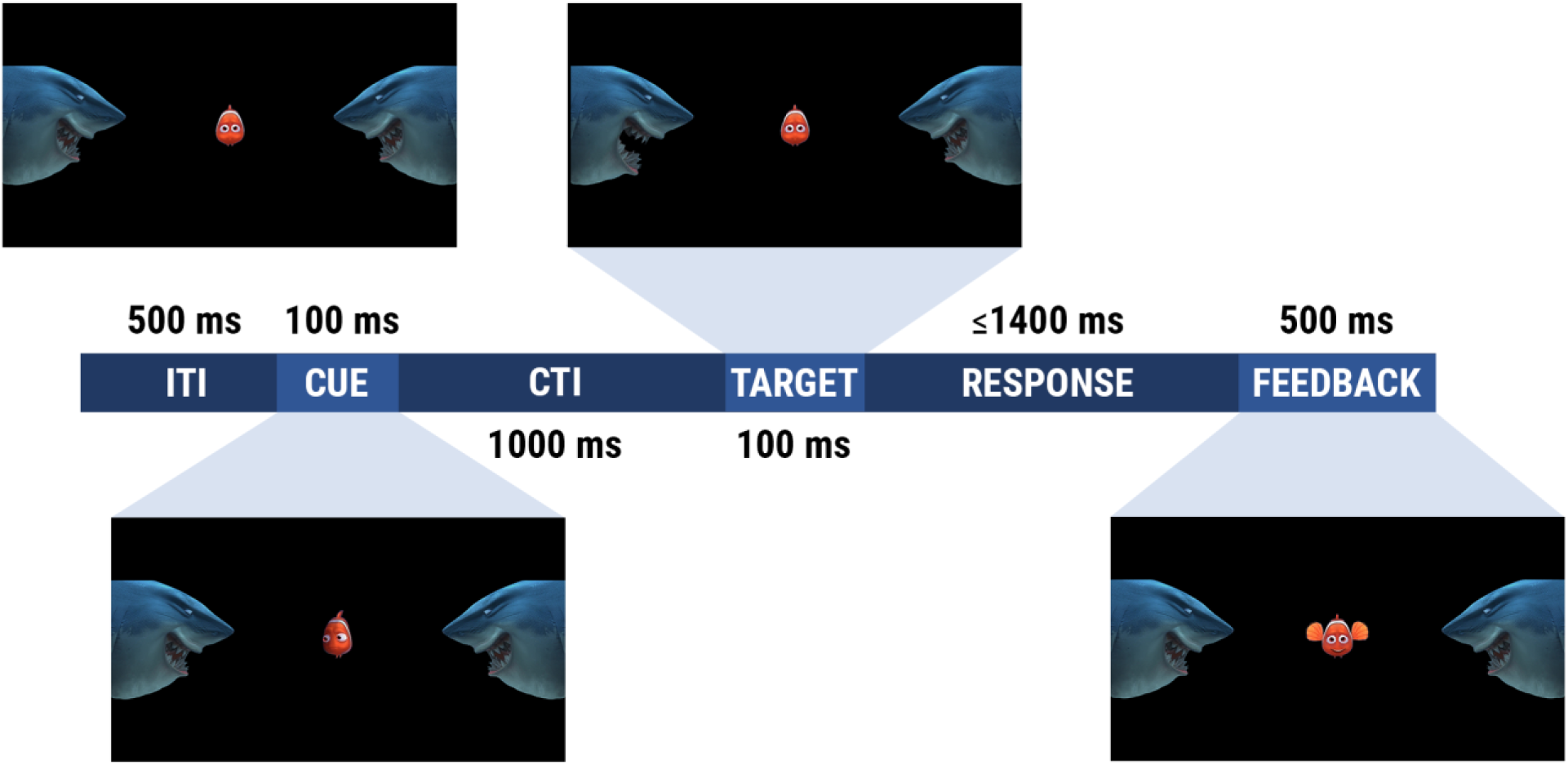
Schematic illustration of the cued covert visuospatial attention task. Participants were instructed to fixate on the centrally placed fish, which would cue participants to attend left or right in anticipation of the target stimulus. After a cue-target interval of 1100 ms, one of the two flanking sharks would open its mouth (target) wider for 100 ms. Participants had up to 1.4 seconds to press the response button corresponding with the direction of the target during validly cued trials only. Pressing the response button would immediately trigger the feedback image before moving on to the next trial. Correct responses triggered a positive feedback image where the central fish appeared to become slightly enlarged and flap its fins, while incorrect responses triggered a negative feedback image which consisted of a fish skeleton.

### Behavioral data analysis

Independent samples t-tests were conducted to examine the reaction time (RT) and sensitivity index (d-prime) between groups. The sensitivity index is derived per signal detection theory, where information from hits and false alarms is used to assess target discrimination (Green and Swets, 1966). Since the task design is a two-alternative forced-choice (2AFC) recognition test, d-prime was corrected by dividing by the square root of two (Macmillan and Creelman, 2005). If a participant did not make any misses, the hit rate was corrected by subtracting 0.5 from the number of hits. Likewise, if a participant did not make any false alarms, the false alarm rate was corrected by adding 0.5 to the number of false alarms. We also conducted post-hoc paired t-tests to examine how RT varied with cue direction within each group.

### Electrophysiological data acquisition

Continuous EEG was recorded from the scalp with 128 Ag-Cl electrodes through the BioSemi ActiveTwo electrode system (BioSemi, Amsterdam, Netherlands), digitized at 512 Hz. With the BioSemi system, the common mode sense (CMS) active electrode and driven right leg (DRL) passive electrode form a feedback loop, which functions as the reference. More information about the referencing and grounding conventions used by the BioSemi active electrode system can be obtained at www.biosemi.com/faq/cms&drl.htm. The EEG data were processed offline using custom in-house scripts for MATLAB R2020a (MathWorks, Natick, MA) and the following MATLAB toolboxes: EEGLAB (Delorme and Makeig, 2004) and FieldTrip toolbox (Oostenveld et al., 2011).

### Data preprocessing

The temporal spectral evolution (TSE) technique was used to characterize alpha-band oscillatory activity during the cue-target interval (CTI) (Salmelin and Hari, 1994; Foxe et al., 1998). The raw EEG data were re-referenced to the average reference for analysis after acquisition. Continuous EEG data were bandpass filtered using a Chebyshev Type II filter with half-amplitude cutoffs of 8 and 14 Hz. Because filtering data with such a narrow bandpass introduces temporal smearing (less than 100 ms), the timing of alpha-related effects is approximate. Bad channels were identified and interpolated. A moving window artifact rejection procedure was applied in which trials were removed if voltage values were above 150 microvolts or at least two standard deviations from the maximum and minimum mean values. After artifact rejection, the data were epoched to 300 ms pre-cue and 1300 ms post-cue and then full-wave rectified. Rectifying the data resulted in all positive-valued data. The rectified epochs were re-baselined to mean voltage over the 100 ms period preceding the cue onset and averaged separately for each cue direction. Thus, voltage values in epoched data represent absolute change in alpha power from baseline levels.

### Data analysis

A grand average waveform was generated for each group for the pre-selected regions-of-interest (ROIs) in the parieto-occipital region of the left and right hemispheres. The left hemisphere ROI is the average of electrode PO3 and the four neighboring electrodes. Likewise, the right hemisphere ROI is the average of electrode PO4 and the four neighboring electrodes. These ROIs were chosen based on the extensive literature on lateralized alpha suppression (Kelly et al., 2010; Murray et al., 2011; Banerjee et al., 2015) (Kelly et al., 2010; Murray et al., 2011; Banerjee et al., 2015) A 4-way mixed ANOVA was performed with the factors of group (deaf, hearing), cue direction (left, right), ROI (left, right), and time window (500-750 ms, 751-1000 ms) on mean alpha-power amplitudes. Pairwise paired t-tests were performed post-hoc across cue left and cue right conditions for activations within each ROI to examine alpha-band spatial attentional modulations within each hemisphere for each group. Scalp topographic maps demonstrating the spatial distribution of alpha power for each cue condition were generated for each group at 750 and 1000 ms.

Next, we wanted to measure topographical differences in alpha-band group-averaged TSE waveforms between groups to examine our hypothesis that general alpha-band related spatial attention deployment might incorporate a more extended occipital-parietal network in deaf individuals. To do this, we collapsed across cue directions between 500 and 900 ms. We quantified topographical differences using global dissimilarity (DISS), which is a reference-free metric and is calculated by taking the root mean square of the differences between two instantaneous GFP-normalized vectors across all electrodes for each time point using a sliding window of 50 ms (Lehmann and Skrandies, 1980; Manly, 1991; Murray et al., 2008). The DISS metric can range from 0 to 2, with 0 indicating topographic equivalence and 2 indicating topographic inversion. To reduce the effects of outlier subjects (Efron and Tibshirani, 1994), DISS was analyzed using a nonparametric Monte Carlo bootstrapping procedure, also colloquially referred to as topographical ANOVA (TANOVA, (Murray et al., 2008). Separate clustering was performed where only clusters meeting or exceeding a *P*-value < 0.05 for at least ten consecutive time samples were considered reliable (Guthrie and Buchwald, 1991). Our bootstrap technique involved resampling the subject DISS values to obtain a distribution of average differences for the null hypothesis that the hearing and deaf groups contrasted were drawn from the same distribution. We randomly sampled 2000 iterations across subjects with replacement to form “test groups” and calculated DISS values. These distributions were used to estimate the probability (*P*-value) that our observed DISS metric could be obtained by chance if our two experimental groups were indeed drawn from the same population. To this end, we established a 95% confidence interval for DISS differences observed between randomly selected “test groups” and tested our observed group differences against this distribution (Efron, 1981).

Additionally, we performed an exploratory post-hoc analysis of alpha activity that appears to occur earlier over the midline parietal region in deaf participants for future hypothesis generation. We created a midline ROI, averaging the EEG activity in electrode A20 and the four neighboring electrodes, and generated a TSE waveform for this ROI (see Figure 5). For statistical testing, cluster-corrected, non-parametric statistical analysis was conducted (see Supplementary Figure S1). For all statistical analyses, alpha criterion of 0.05 was used and corrected for multiple comparisons when appropriate.

## Results

### Behavioral performance

All participants completed both blocks of the experiment, consisting of a total of 368 trials. For d-prime, deaf participants were more accurate than hearing participants (*t*_19_ = 2.06, *P* = 0.046, Cohen’s *d* = 0.64). The d-prime was 2.68 ± 0.68 for hearing participants and 3.06 ± 0.49 for deaf participants (Figure 2). Deaf participants (423.1 ± 70.7 ms) also responded more quickly to validly cued targets compared to hearing participants (494.8 ± 57.2 ms), resulting in a statistically significant difference between groups; *t*_19_ = 3.53, *P* = 0.001, Cohen’s *d* = 1.11 (Figure 2). The finding that deaf native signers responded significantly more quickly is in line with previous research (Neville and Lawson, 1987a; Loke and Song, 1991; Chen et al., 2006; Nava et al., 2008; Bottari et al., 2010). Post-hoc analysis of RTs separated by cue direction suggested a right-field advantage for both groups. For the deaf group, RT was 432 ± 72.5 ms for left cue trials and 414.4 ± 70.9 ms for right cue trials; *t*_19_ = 3.43, *P* = 0.0028, Cohen’s *d* = 0.25. For the hearing group, RT was 504.53 ± 60 ms for left cue trials and 485.03 ± 57 ms for right cue trials; *t*_19_ = 3.43, *P* = 0.0028, Cohen’s *d* = 0.33.

**Figure 2.**
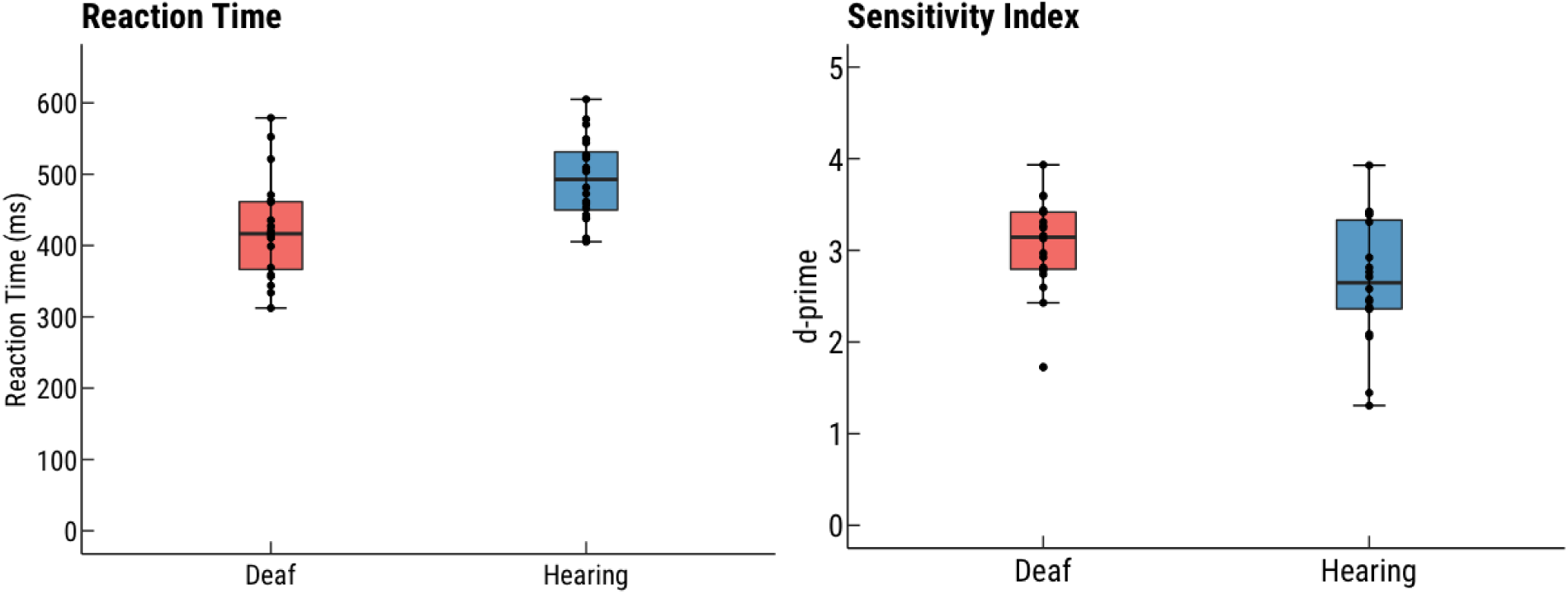
Box-and-whiskers plots (with over-plotted individual data points) for reaction time and sensitivity index (d’) for deaf and hearing participants. Deaf participants were significantly faster (t_19_ = 3.53, P = 0.001; Cohen’s d = 1.11) and more accurate (t_19_ = 2.06, P = 0.046; Cohen’s d = 0.64) than hearing participants.

### Spatially-specific alpha-band activity

The alpha-band TSE waveform for left and right cue trials in the parieto-occipital ROIs over the left and right hemispheres of hearing and deaf participants can be observed in Figure 3. Results from the 4-way mixed ANOVA showed main effects of hemisphere (*F*_1,38_ = 17.36, *P* < 0.001, η^2^_*p*_ = 0.31). There was also a significant interaction effect for hemisphere x cue direction (*F*_1,38_ = 31.62, *P* < 0.0001, η^2^_*p*_ = 0.45), hemisphere x time (*F*_1,38_ = 6.55, *P* = 0.015, η^2^_*p*_ = 0.15), and for group x time (*F*_1,38_ = 4.89, *P* = 0.033, η^2^_*p*_ = 0.11).

**Figure 3.**
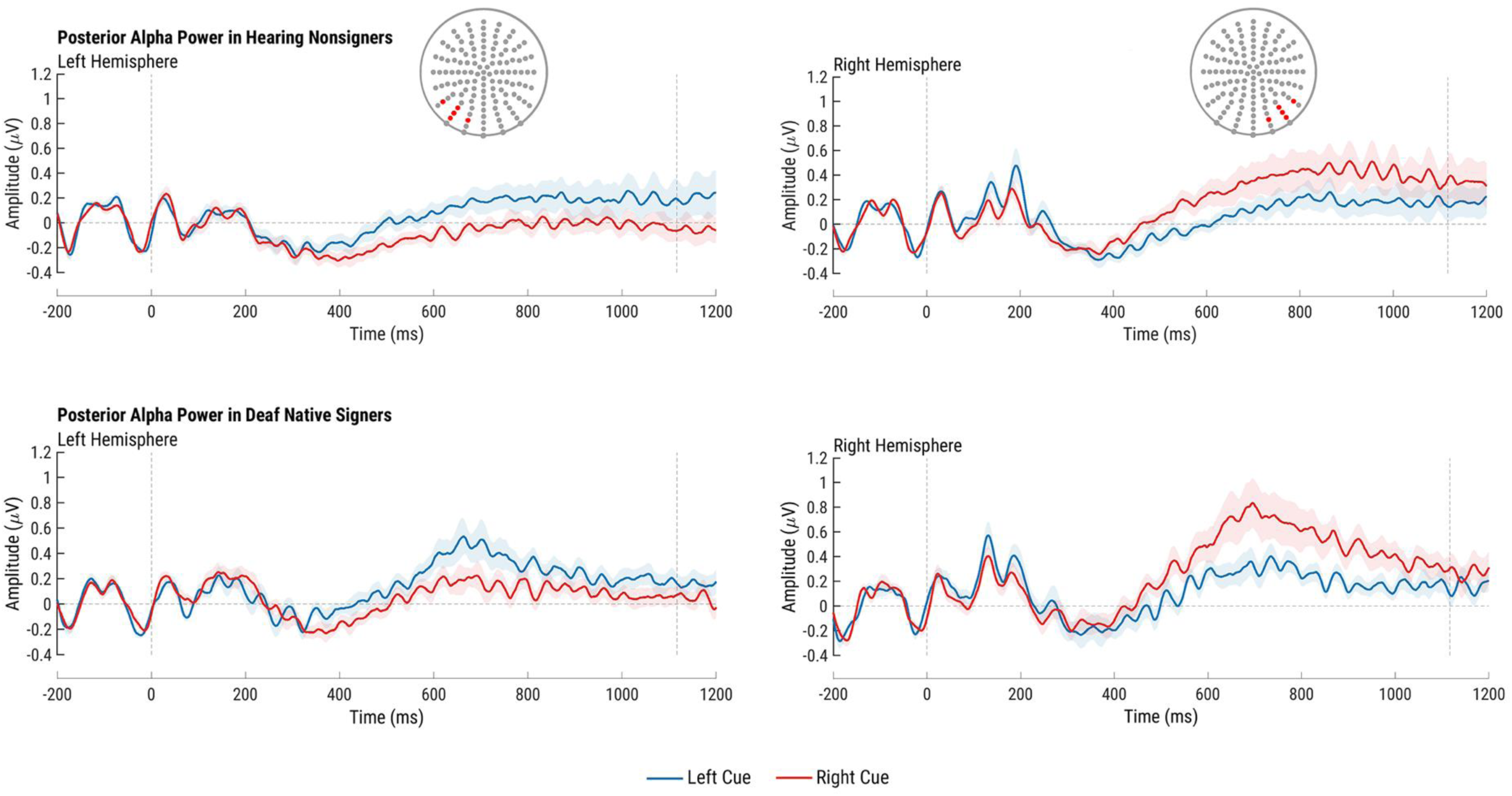
Alpha-band (8-14 Hz) activity over the left and right parieto-occipital ROIs for hearing non-signers (top row) and deaf native signers (bottom row) during the interval between the cue (0 ms) and the target (1100 ms). Higher alpha power was observed over each posterior ROI for both groups when the ipsilateral hemifield was cued.

It is not clear whether the interaction of hemisphere x cue direction is based on a contralateral increase in alpha-band activity, given that there is higher alpha power over the right hemisphere compared to the left, which is in line with previous work (Foxe et al., 1998; Fu et al., 2001; Banerjee et al., 2011). To confirm that the increase in alpha power is lateralized to the unattended hemifield, we conducted pairwise paired t-tests for each group comparing cue conditions within each hemisphere. Higher alpha power was observed for the cue left vs. cue right condition over the left hemisphere in hearing participants (*t*_39_ = 5.95, *P* < 0.0001, Cohen’s *d* = 0.94) and deaf participants (*t*_39_ = 5.67, *P* < 0.001, Cohen’s *d* = 0.90). Similarly, significantly higher alpha power was observed for the cue right vs. cue left condition over the right hemisphere in hearing participants (*t*_39_ = 3.54, *P* = 0.001, Cohen’s *d* = 0.56) and deaf participants (*t*_39_ = 3.56, *P* = 0.001, Cohen’s *d* = 0.56). The lateralization of alpha-band activity over the parietal region can be observed in Figure 4, which shows the topographic distribution of instantaneous alpha-band oscillatory activity at 750 and 1000 ms post-cue for hearing and deaf participants. There is greater activity over the parieto-occipital region ipsilateral to the cued hemifield at both time points. In line with previous evidence of a parietal asymmetry in attentional control favoring the right parietal lobe (Gitelman et al., 1999), there is greater activity over the right parietal region for both groups relative to the left.

**Figure 4.**
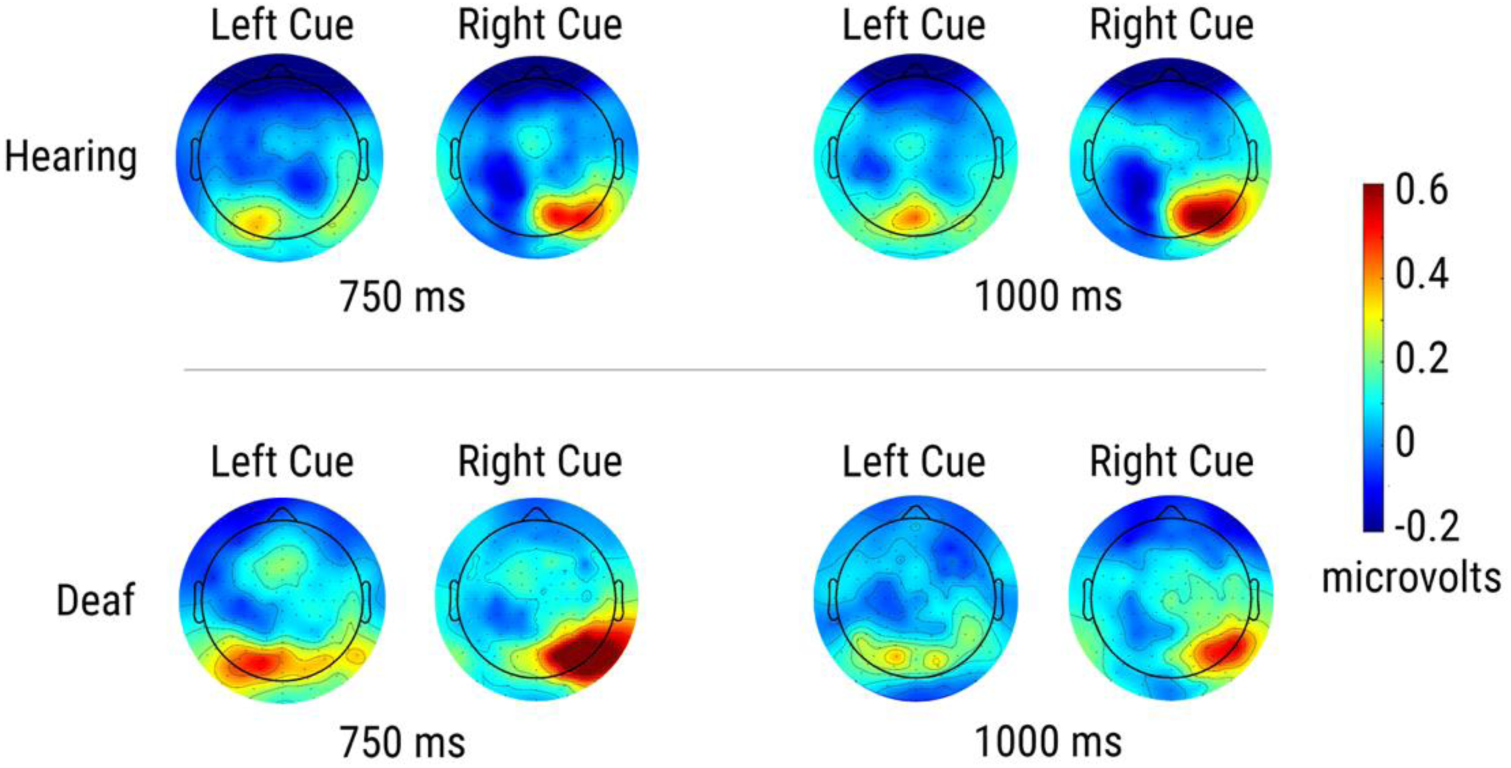
Scalp topographic maps for average alpha power at 750 and 1000 ms post-cue for left and right cue trials in hearing non-signers (top row) and deaf native signers (bottom row). Distinct topographical patterns can be observed over the parietal ROIs corresponding with the cued direction for both groups.

Interestingly, pairwise paired t-tests comparing groups by time frame found that deaf participants exhibited greater alpha activity during 500-750 ms compared to hearing participants (*t*_79_ = 3.64, *P* < 0.001, Cohen’s *d* = 0.41), but no difference was found between groups during the 751-1000 ms time window. This finding suggests that the onset of alpha activity occurs earlier in deaf individuals.

### Right parietal control of spatial attention

The second part of our study explored the possibility of sensory-substitution processes in the deployment of attention in deaf individuals. We hypothesized that the distribution of alpha-band activity over parietal scalp would be greater in the anticipatory period in deaf individuals, reflecting an expansion/sequestration of attentional control systems to sensory-specific regions that were redundant (i.e., unused auditory control systems). Figure 5 shows the alpha-band scalp topographies with cue conditions collapsed over 100 ms intervals between 500 and 900 ms during the cue-target interval. Hearing participants showed a highly focused right parieto-occipital alpha distribution in the midline around 600 ms post-cue that became more distributed over the parieto-occipital region during the second half of the cue-target interval. There was a highly focused alpha-band distribution in the middle parietal region in deaf participants present at 500 ms post-cue that quickly became more widely distributed over the right parietal region. Figure 6 shows the results of the bootstrapped TANOVA of the DISS values calculated between each group for each time point during the epoch, demonstrating that the topographical maps for each group were significantly dissimilar (*P* < 0.05) around 500-850 ms post-cue.

**Figure 5.**
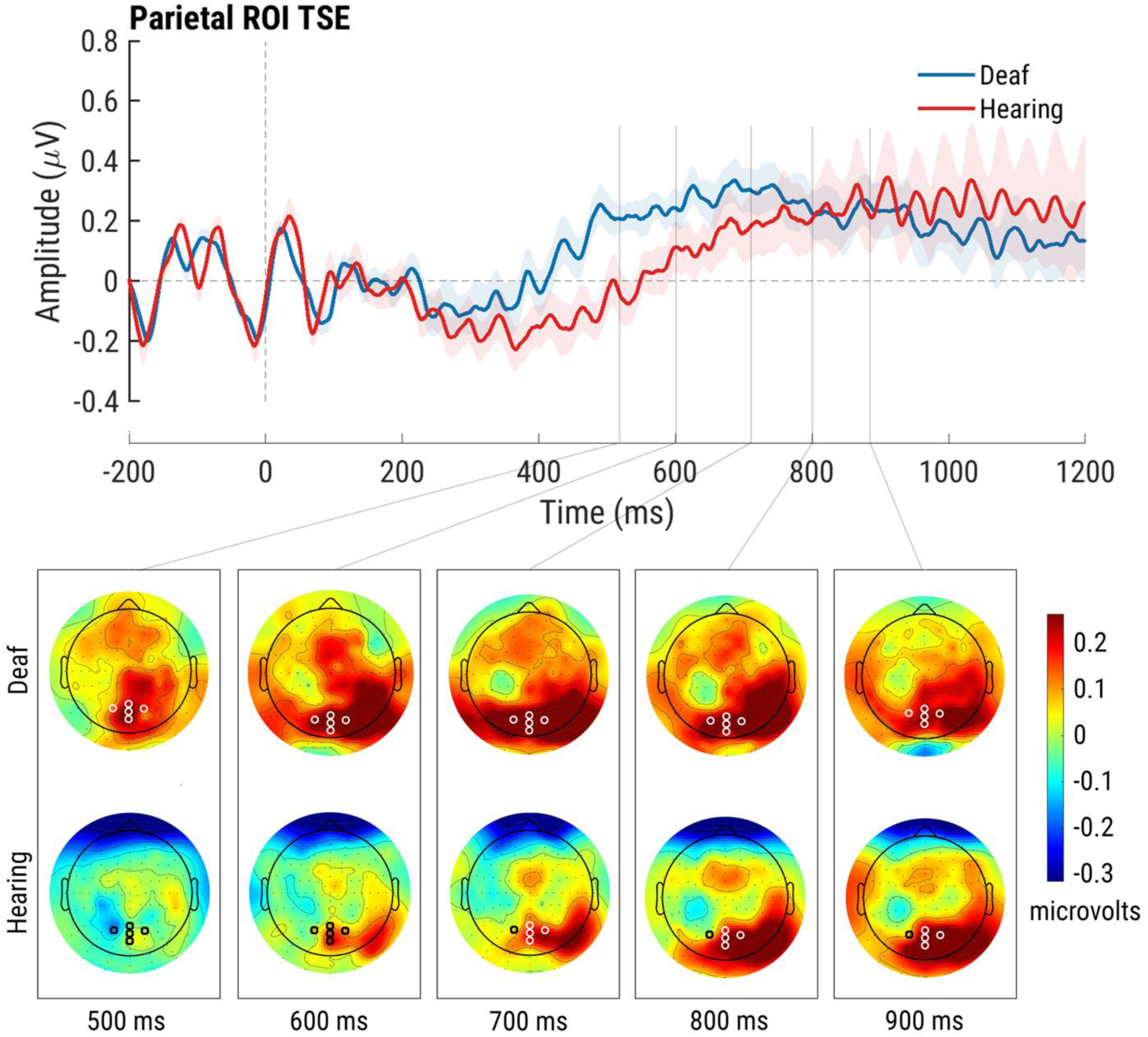
Alpha-band activity over the midline parietal ROI with cues collapsed (i.e., left plus right cues). Corresponding scalp topographies are shown at intervals of 100 ms for the time points between 500 and 900 ms post-cue.

**Figure 6.**
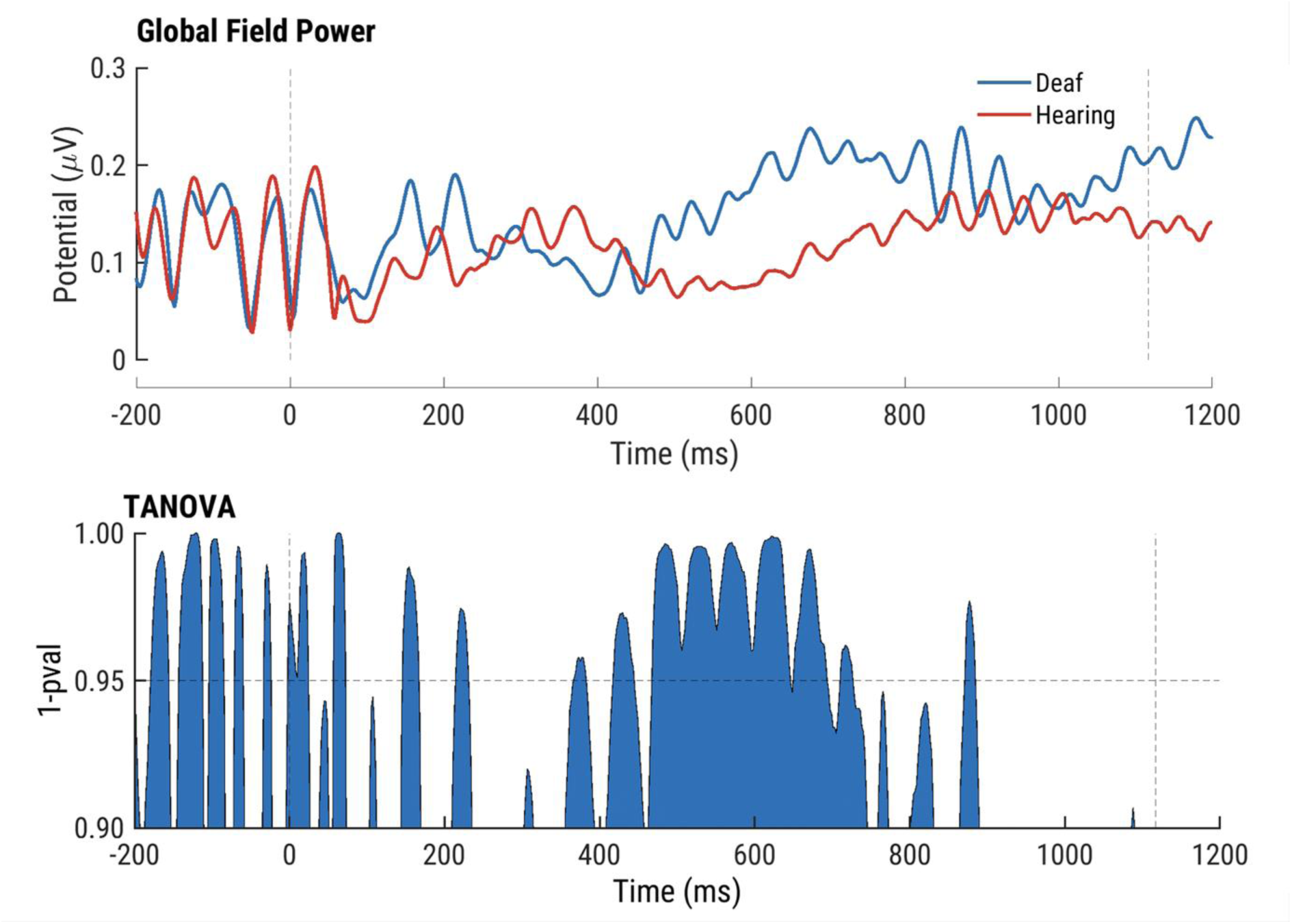
Global field power (GFP) and the results of the analysis of topographical dissimilarities (TANOVA) between deaf and hearing participants over the time course of the anticipatory epoch (the cue-target interval). Topographical distributions were significantly dissimilar during the 400-700 ms post-cue period and converged around 900 ms.

### Exploratory analysis of earlier onset midline posterior alpha activity in deaf individuals

With cue conditions collapsed and based on post-hoc visual inspection of the data and the group x time interaction noted in the main ANOVA analysis above, we observed an apparent increase in alpha-band activity that occurred earlier in deaf participants over the midline parietal region. This activity was already present in deaf participants at 500 ms but not in hearing participants (Figure 5). The difference in alpha power between groups over the midline ROI can be appreciated further in Figure 6, where alpha power appears to be elevated in deaf participants starting around 300 ms and continuing until 800 ms. In addition, post-hoc exploratory statistical cluster plots revealed earlier alpha activity over the parietal region in deaf participants between 500-800 ms (see Figure S1), an effect we had not specifically predicted.

## Discussion

Here, we set out to test whether deaf individuals would show increased covert spatial attention performance in the visual modality, whether the deployment of visuospatial attention would be associated with lateralized alpha-band oscillatory neural mechanisms that have been repeatedly identified in the hearing population, and if these anticipatory alpha-band processes would show evidence for extended sources within the parietal cortex that would suggest the recruitment of additional non-visual (i.e. auditory) attentional control regions, a finding that would support the sensory-substitution hypothesis. The data were, in part, consistent with each of these three predictions. When considering performance on the covert spatial attention task, deaf individuals were more accurate in identifying cued targets and considerably quicker in responding to these targets, demonstrating clearly better performance on this task. Considering the second prediction, deaf individuals showed typical patterns of lateralization of alpha-band anticipatory oscillatory activity in the period between the spatial cue and the subsequent target stimulus, mimicking in large part the effects seen in hearing controls. It should be noted, however, that this lateralized alpha-band activity was not of greater amplitude in deaf individuals. Considering the third prediction, topographic analysis of the anticipatory alpha-band activity, without regard for the directionality of attention (i.e., with left and right cueing conditions collapsed), showed clearly increased extent of the distribution of alpha over parietal scalp, supporting the notion that additional parietal regions were recruited in the execution of this task in deaf individuals. Lastly, although not specifically hypothesized, both the main ANOVA, which included time as a factor, and *post-hoc* analysis, showed that alpha-band oscillatory activity was deployed markedly earlier during the cue-target interval in deaf individuals, which may partially explain the faster response times and improved accuracy in this group. These findings are discussed in more detail in what follows.

### Enhanced visuo-spatial task performance in deaf participants

We observed a statistically significant difference in reaction times between groups with deaf participants responding considerably more quickly than hearing participants. This substantial speeding of responses did not come, however, at the expense of accuracy, as deaf individuals also proved more accurate at detecting targets than hearing individuals. Although improved visuo-spatial attention abilities have not been an entirely consistent finding in deafness (see Dye and Bavelier, 2013 for a review; Holmer et al., 2020), other research groups have shown similar speeding of responses to peripheral targets in deaf individuals (Loke and Song, 1991; Codina et al., 2011, 2017), and it is perhaps intuitive that individuals who must rely more heavily on one sensory system for spatial alerting, and spatial information processing, would evince better tuned performance in that sensory system. That additional “unused” cortical regions might be recruited in the service of this compensatory mechanism would also align with this improved ability. There is certainly evidence in the literature for this type of cortical cross-sensory recruitment in individuals who have lost a sensory system, be it vision or audition (Nishimura et al., 1999; MacSweeney et al., 2002; Striem-Amit et al., 2012), and as will be discussed below, topographic analysis here points to engagement of a more extensive tract of parietal cortex by the deaf participants in this study.

It is also worth noting that there appears to be a right-field advantage whereby both groups responded more quickly on right cue trials compared to left cue trials. This right-field advantage could be due in part to the so-called Poffenberger effect, where reaction times are found to be slower when the motor response (all participants responded using their right hand) and the visual field condition are contralateral (requiring interhemispheric communication) compared to when the motor response and the visual field condition are ipsilateral (requiring intrahemispheric transfer) (Poffenberger, 1912; Berlucchi et al., 1977; Clarke and Zaidel, 1989; Saron et al., 2003a, 2003b). However, Parasnis and Samar (1985) also saw an overall right-field advantage across all conditions when a bimanual response mode was used during a cued visuospatial task, which they attributed to a “structurally determined attentional bias toward the right side of space” independent of more specific stimulus processing mechanisms.

### Evidence of spatially-specific alpha-band activity

Deaf native signers and hearing non-signers alike exhibited clear alpha-band lateralization over parieto-occipital scalp corresponding to the direction of their covert spatial attention, replicating a well-established finding in the literature (Worden et al., 2000; Yamagishi et al., 2003; Kelly et al., 2005, 2006, 2009; Sauseng et al., 2005; Rihs et al., 2007, 2009). According to the alpha suppression hypothesis (Foxe and Snyder, 2011; Van Diepen et al., 2019), the pattern of alpha lateralization observed in deaf native signers suggests that they are fully capable of selectively inhibiting irrelevant visual space in anticipation of the appearance of target stimuli in the cued visual hemifield. In other words, deaf native signers do not appear to be continuously monitoring the periphery, but rather, are deploying attention in a phasic trial-by-trial manner in response to environmental cues, at least under the current task parameters.

This finding is interesting when you consider the findings from Proksch and Bavelier (2002) that deaf native signers appeared to be more sensitive to peripheral distractors than hearing participants. In that study, deaf native signers, hearing native signers, and hearing non-signers were instructed to perform a search task while ignoring distractors that appeared in the center or the periphery. Unlike both hearing groups, who were more distracted by central distractors, the deaf participants were more distracted by peripheral distractors, suggesting that deaf native signers deploy more attentional resources to continuously monitor the periphery. Also, another study by Bosworth and Dobkins (2002) found that deaf participants performed better when the target stimulus was presented in the periphery among distractors than when it was presented alone. Importantly, this effect was observed when the duration of the cue-target interval was 600 ms but not when it was 200 ms. To explain this finding, the authors proposed two possible interpretations: (1) ignoring distractors requires increased top-down cognitive effort, which improves processing of the target stimulus; and (2) sensory bottom-up processing of the target is enhanced by the presence of distractors surrounding the target stimulus. The former interpretation appears to be more likely when one considers that the effect was only observed for the 600 ms interval and that top-down processing requires more time to take effect. Our findings may lend support to this interpretation given that alpha activity in the posterior parietal cortex begins earlier in deaf participants. With increased top-down cognitive effort to suppress peripheral distractors over a longer period, deaf participants may be primed to process the target stimulus faster than hearing participants. In the current study, it should be noted that irrelevant targets were never to be responded to, so the task promotes selective suppression of these potentially distracting inputs.

### Potential sensory substitution processes in attentional control

In previous work comparing alpha-band activity during audio- and visuo-spatial tasks, we observed a highly focused alpha distribution over the right parieto-occipital region shared by both sensory modalities at 600-700 ms post-cue, with cue conditions collapsed (Banerjee et al., 2011). Immediately thereafter, we observed the emergence of two highly distinct but adjacent foci of alpha distribution associated with each sensory modality. These findings suggest that there is an interaction between supramodal and sensory-specific spatial attention mechanisms in the parietal cortex during the anticipatory deployment of spatial attention. When comparing alpha topographies between deaf and hearing participants with cue conditions collapsed, we saw a midline focus appearing much earlier in deaf participants that became more broadly distributed over the parieto-occipital region, especially between 500-800 ms post-cue. Exploratory analyses provided further support that the enhancement of alpha activity occurs earlier in deaf participants.

This finding implicates the engagement of a more extensive network in the parietal attention control system in deaf individuals, which may explain the enhancement in visuospatial attention abilities in deaf native signers, something that will need to be explicitly tested in future work. The potential repurposing of unused cortical regions typically associated with the modulation of audio-spatial attention may also explain the findings reported in Bonacci et al. (2019), where alpha lateralization as a correlate of audio-spatial attention was explored. In participants with bilateral symmetric sensorineural hearing loss (at least 25 dB loss in the better ear), there was no significant alpha lateralization, unlike in hearing participants. Limited exposure to sound from birth may lead to reorganization of the attentional control region in the posterior parietal cortex where the function (i.e., attentional control) is preserved but adapted to process input from a different sense (i.e., vision)—a concept termed *functional preservation* (Cardin et al., 2020). It may be worth exploring whether the possibility of a visual “takeover” of the region typically used for audio-spatial attentional control makes it difficult to process auditory inputs after reorganization has already taken place. Of note, Bonacci and colleagues did not report the use of sign language and hearing aids in their study population. Data on hearing aid use would be critical as it has been reported that sound localization is worse when wearing hearing aids, potentially interfering with the development of audio-spatial attentional skills in this population (Van den Bogaert et al., 2006).

### Further limitations and considerations

It is worth mentioning that alternate exogenous (bottom-up) attentional capture mechanisms may also be at play here in this ostensibly endogenous voluntary attentional deployment task. Abrupt changes in the appearance of the target in the periphery may have triggered more automatic exogenous orienting of attention and Bottari and colleagues (2008) have suggested that the peripheral visual enhancement observed in deaf people may rely on this exogenous component of visual attention. In their study, no difference in performance between deaf and hearing participants was observed when the exogenous component of attentional orienting was removed by triggering a global transient instead of a local transient in the periphery during a change blindness paradigm (Bottari et al., 2008). Thus, the deaf participants in our study, in addition to successfully endogenously biasing their spatial attention in anticipation of the impending target, may have received an additional bump in performance due to increased efficiency in exogenously orienting attention to the explicit arrival of the target stimuli. Of course, this aspect of the study was not explicitly manipulated here, so no firm conclusions can be drawn from the current results.

At first glance, our findings are also not completely consistent with the results of another visuospatial cueing study where it was shown that deaf native signers responded more quickly to invalidly cued trials but not to validly cued trials (Parasnis and Samar, 1985). Paradigm differences are the likely source of differences here, since we did not require our participants to respond to invalidly cued targets, so endogenous attention could be directed wholly at the impending location of a potential valid target without the need to also partially monitor invalidly cued space. It is likely that the divided attention required to respond to both validly and invalidly cued targets substantially reduces the amount of spatial biasing that can be deployed under such circumstances. It is also possible that the increased exogenous attentional capture seen in deafness in the Bottari study discussed above, could explain the increases in responding to uncued targets shown in the Parasnis and Samar study.

Another design feature of the current study that is not entirely consistent with the paradigms that have been typically used to assess endogenous visuospatial attention deployments, was the use of a “gaze-cue” to direct spatial attention rather than the more typically used symbolic arrow cues. That is, we used changes in the directionality of the eyes of a centrally presented cartoon fish to indicate to which side of space covert attention was to be directed. Friesen and Kingstone (1998) found that, like peripherally-located symbolic cues, centrally-located gaze cues facilitated exogenous, or reflexive, shifts of covert attention, when the cue-target interval was short (e.g., as early as ∼100 ms post-cue). In other words, attention was reflexively drawn towards the cued location resulting in faster RTs during validly cued trials compared to invalidly cued trials – a phenomenon called the gaze-cueing effect (GCE). However, the GCE was only observed when the cue-target interval was short (e.g., as early as ∼100 ms post-cue) but not when the cue-target interval was long (e.g., ∼1,000 ms). Nevertheless, there is some evidence that attentional shifts facilitated by gaze cues are activated by different neural systems (Hietanen et al., 2006, 2008; Frischen et al., 2007). When participants are expected to shift attention to the direction of a gaze cue, there is greater activity in the superior temporal sulcus (STS), an area involved in face and gaze processing, compared to when arrow cues are used (Hoffman and Haxby, 2000; Hooker et al., 2003). This finding is especially interesting when you consider that deaf individuals show greater recruitment of the posterior STS when processing visual stimuli (Bavelier et al., 2001).

One study comparing the effect of gaze cues and arrow cues between deaf signers, hearing signers, and hearing non-signers found no evidence of increased GCE in deaf signers, but a comparable arrow-cueing effect was observed across groups (Heimler *et al*., 2015). It may be worth exploring the effect of arrow cues on alpha modulations in deaf native signers in a variant of the current paradigm.

Also, several recent studies have questioned whether lateralized alpha amplitude modulation has a causal role in visuospatial attention control (Keitel et al., 2019; Antonov et al., 2020; see Peylo et al., 2021 for a review). Antonov and colleagues (2020) reported that an increase in steady-state visual evoked potential (SSVEP) amplitudes preceded lateralized suppression of alpha activity, which suggests that alpha modulation is instead the consequence of shifted attention. In contrast, Gundlach et al. (2020) reported the opposite pattern. This contradiction in timing effects may be explained by temporal and spectral smearing caused by the use of a narrow filter. Also, several studies using noninvasive brain stimulation approaches such as transcranial magnetic stimulation (TMS) or transcranial electrical stimulation (tES) provide evidence of a causal role of alpha modulation in visuospatial attention control via entrainment of alpha activity (Capotosto et al., 2009; Romei et al., 2010; Sauseng et al., 2011; Kasten et al., 2020), and in prior work from our research group, we showed that target discriminability and response times were significantly predicted on a trial-by-trial basis by the strength of lateralized alpha-band amplitude in the 500 ms preceding the target (Kelly et al., 2009).

What is not yet clear is whether early sign language exposure has an independent effect on preparatory attentional processes. Deaf native signers are considered ideal language models because they acquire sign language vertically from their deaf parents and achieve typical developmental language milestones, potentially earlier than hearing children (Bonvillian et al., 1983; Petitto et al., 2001). Thus, it is logical to consider that the early and daily use of a signed language could play a role in the modulation of visuospatial attention, primarily because native signers typically focus on or near the eyes of the interlocutor while covertly attending to their hands and arms moving outside of the foveal space (Emmorey et al., 2009). One way to separate the possible impact of early sign language acquisition on visuospatial attention would be to introduce hearing native signers as a third group. However, there is evidence that hearing native signers do not display the peripheral visual advantages seen in deaf individuals, suggesting that these advantages are due to early deafness and not the early acquisition of a visual language (Neville and Lawson, 1987b; Bavelier et al., 2001; Bosworth and Dobkins, 2002; Proksch and Bavelier, 2002; Dye et al., 2009; Heimler et al., 2015).

There are a number of studies that report visuospatial attentional deficits and reduced cognitive control in the deaf population, mostly in children (Myklebust and Brutten, 1953; Quittner et al., 1990, 1994; Mitchell and Quittner, 1996; Rettenbach et al., 1999; Rothpletz et al., 2003; Sladen et al., 2005; Chen et al., 2010). Some suggest that poorer performance on attentional and cognitive tasks among deaf people can be attributable to the auditory deficit (Myklebust and Johnson, 1964; Furth, 1966; Mitchell and Quittner, 1996; Conway et al., 2009; Kronenberger et al., 2014). The perceptual deficit hypothesis proposes that a “substantial deficit in one sensory modality could affect the development and organization of the other sensory systems’’ (Turkewitz and Kenny, 1982; Radell and Gottlieb, 1992; Pavani and Bottari, 2012). In contrast to the perceptual deficit hypothesis, the compensatory hypothesis proposes that the loss of a sense can enhance the remaining senses due to a greater reliance on these senses (Neville, 1990; Grafman, 2000; Pavani and Bottari, 2012). Our findings suggest that the brain areas typically reserved for the processing of audiospatial inputs may be functionally reallocated in deaf native signers, which is made evident by the earlier and broader distribution of posterior alpha activity compared to hearing participants.

Some argue that the studies reporting attentional deficits in deaf people are due to the heterogeneity of the subject group and the failure to control for differences in etiology of deafness, age of onset, and preferred language or mode of communication (Hoemann, 1978; Bavelier et al., 2006; Dye and Bavelier, 2013). Studies have shown that withholding sign language from deaf children who fail to meet spoken language milestones during early childhood puts them at risk of language deprivation syndrome, which can have long-lasting effects on cognition and behavior (Hall, 2017; Hall et al., 2017a, 2017b). Additionally, there is evidence of neuroanatomical differences between deaf native users of ASL and deaf non-native signers (Olulade *et al*., 2014). Interestingly, Heimler et al. (2015) found that, like hearing native signers and hearing non-signers, deaf participants who acquired Italian Sign Language later in life showed a greater gaze-cueing effect in response times compared to those who acquired sign language earlier but only for trials with shorter stimulus-onset asynchrony (250 ms). It is unknown whether delayed language access in the face of deafness, whether spoken or signed, would result in impaired alpha suppression. In order to answer that question, this task will need to be conducted with deaf, non-native signers who did not acquire a signed language until after the critical period of language learning had narrowed— around the age of 5 (Lenneberg, 1967; Mayberry and Eichen, 1991; Hall et al., 2017b). The answer to this question would be very informative given that roughly 95% of deaf children are born to hearing parents, and most of these children do not have the opportunity to vertically acquire a signed language from their parents at home (Mitchell and Karchmer, 2004).

## Conclusion

Assessing the modulation of alpha-band activity during the cue-target interval of a cued covert visuospatial attention task using EEG is a useful approach to understanding how neuroplastic changes associated with deafness affect the ability to modulate the processing of visuospatial information, both relevant inputs and those that should be ignored/suppressed. Impaired anticipatory alpha-band modulation may have clinical relevance as there is a correlation with impaired performance on attention tasks in individuals on the autism spectrum (Murphy et al., 2014), as well as children and adults with attentional deficit hyperactivity disorder (ter Huurne et al., 2013; Vollebregt et al., 2016). However, deaf native signers demonstrated enhanced, not impaired, performance on the current visuospatial attention task. Additionally, they demonstrated lateralization of alpha-band oscillatory neural mechanisms wholly similar to those observed in the hearing population, except that the onset of alpha activity appeared to take place earlier. Lastly, we provide evidence suggesting that additional non-visual attentional control regions are recruited because of limited access to sound in this population. Our findings help provide further insight regarding the parietal attentional control system in deaf native signers that may provide more context to some conflicting findings in the existing literature.

## Author Contributions

IAD, JJF, and EGF conceived of the study and regularly consulted during the development of the analyses and data representations. Data collection and recruitment were led by IAD with significant assistance from JX and MMS. IAD performed the primary data analyses and statistical tests in close consultation with JJF, EGF, and KDP, created the illustrations, and wrote the first substantial draft of the manuscript. JJF and EGF provided substantial editorial input to IAD during multiple drafts of the manuscript. All authors read and approved the final version for publication, attest to the accuracy of the work reported, and agree to be fully accountable for all aspects of the work.

## Acknowledgments

The authors thank the entire team at The Frederick J. and Marion A. Schindler Cognitive Neurophysiology Laboratory at the University of Rochester for their continued support of the research efforts of our group. We also thank our participants for donating their valuable time and effort to this study.

## Abbreviations List

PPC: Posterior Parietal Cortex
ASL: American Sign Language
RT: reaction time
EEG: electroencephalography
TSE: temporal spectral evolution
CTI: cue-target interval
ROI: region of interest
DISS: global dissimilarity
GFP: global field power
GCE: gaze-cueing effect
ANOVA: analysis of variance
TANOVA: topographical analysis of variance

## Supplemental Figures

**Figure S1.**
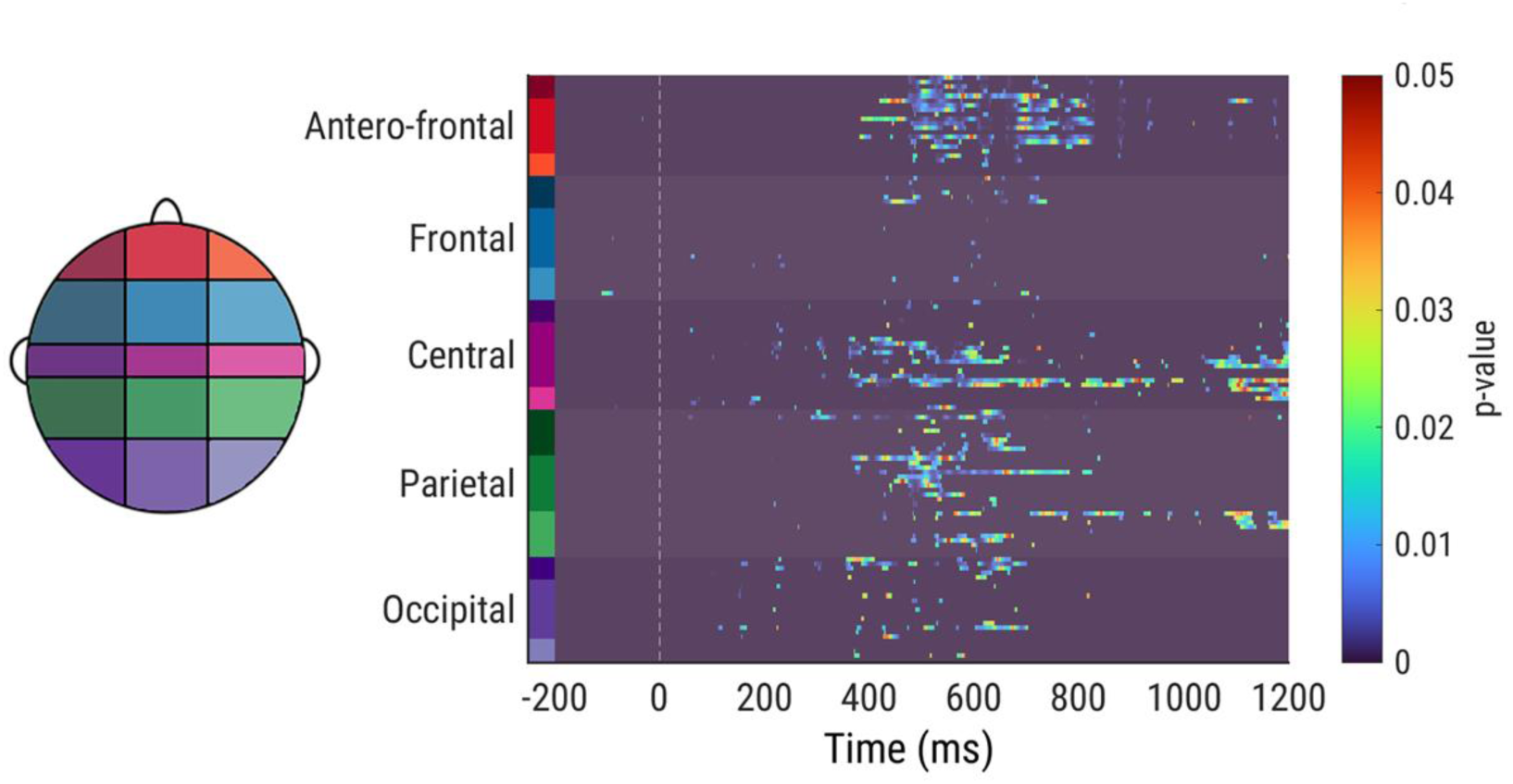
Cluster-corrected plot of significant values obtained from independent t-tests between deaf and hearing participants for all electrodes and time points during the cue-target interval. Much of the significant differences were observed between 400 and 800 ms post-cue especially over the parietal and antero-frontal regions.

